# Clubroot disease in soil: An examination of its occurrence in chemical and organic environments

**DOI:** 10.1101/2024.05.29.596495

**Authors:** Zakirul Islam, Quoc Thinh Tran, Motoki Kubo

## Abstract

Clubroot is a disease in cruciferous plants caused by the soil-borne pathogen *Plasmodiophora brassicae*. This pathogen rapidly spreads in soil, and plant growth is inhibited by infection with spores. In this study, the development of clubroot disease in *Brassica rapa* var. perviridis was investigated in different soil environments (chemical and organic soils). The bacterial biomass and diversity of the soil and roots were also analyzed in both chemical and organic soils. Bacterial biomass and diversity in the organic soil were higher than those in the chemical soil. The disease severity of plants cultivated in organic soil was lower than that in chemical soil. The number of endophytic bacteria in the roots decreased when the plants were infected with *P. brassicae* in both soil types. Higher bacterial biomass in the soils and roots appeared to reduce the infection of *P. brassicae*.

**Importanc:** Clubroot is a major disease of cruciferous plants caused by soil borne pathogen *Plasmodiophora brassica.* Pesticides are applied in the soil during cultivation for the inhibition of clubroot disease, which demise soil microbiome. Therefore, eco-friendly control methods are required for sustainable cultivation. Soil management is an important strategy to inhibit clubroot disease severity. In this study, the occurrence of clubroot disease in *B.rapa* were compared in two distinct soil environments (chemical and organic). Results show infection rate increased with the increase of spore number for both chemical and organic soil environments. However, the diseases severity was lower in plants cultivated in organic soil. The higher endophytic bacterial biomass and diversity seems to be linked for the disease inhibition in organic plants. These findings suggest a sustainable soil management process to reduce clubroot disease prevalence. Application of organic fertilizers in commented soils could substantially reduce clubroot disease severity in the plants.

## 1. Introduction

Chemical fertilizers have been extensively used in agricultural practices throughout the 20^th^ century to achieve high crop yields [1]. Inorganic forms of chemical fertilizers, particularly those containing nitrogen, phosphorus, and potassium, can be directly applied to the soil and immediately absorbed by plants. Fertilizer application has been responsible for at least a 50% increase in crop yield in the 20th century [2, 3]. However, the continuous application of chemical fertilizers over many years has altered the soil environment [4]. Soil organic matter (SOM), a key attribute of soil health, gradually decreases with the application of chemical fertilizers. In addition, soil microorganisms, which play diverse ecological roles, such as organic compound mineralization, carbon sequestration, and disease suppression, are adversely impacted by both the direct and indirect application of chemical fertilizers [5, 6, 7, 8]. Poor soil quality can lead to persistent crop difficulties and diseases caused by soil borne pathogens [9, 10].

Pathogenic microorganisms cause plant diseases, resulting in considerable losses in crop production. Pathogenic fungi, such as rust and powdery mildew, infect plants from their aboveground components (stems and leaves) and are transmitted through the air [11]. In contrast, root-knot fungi typically reside in the soil for extended periods of time and are transmitted via water within the roots [12]. *Plasmodiophora brassicae*, an obligate plant parasite, can remain dormant in the soil for up to 20 years [13, 14]. It germinates in the roots of cruciferous plants, causing clubroot disease [15]. Infected roots are incapable of absorbing water and nutrients from the soil, resulting in delayed growth and, in certain instances, plant death. Clubroot disease causes a 10–15% loss in total yield worldwide [16]. Fungicides are typically applied to soil during cultivation to reduce the occurrence of clubroot disease; however, suppression practices applying agrochemicals also result in the demise of beneficial microorganisms [17]. Conversely, some studies have found that the prevalence of naturally occurring soil microorganisms can inhibit the development of clubroot disease [18]. Considering eco-friendly cultivation practices, strategies that prioritize soil management with a high level of microorganisms may serve as a sustainable solution for effectively inhibiting clubroot disease in plants.

The soil fertility index (SOFIX) was developed to evaluate soil fertility and determine suitable soil conditions for organic agriculture [19]. This technology enables the preservation and diversification of soil microorganisms by utilizing biomass resources. It has been applied to promote sustainable organic farming. Current research is focused on studying the antagonistic suppression effect of soil-borne pathogens by enhancing soil microbiomes utilizing SOFIX technology. In this study, we examined the onset of clubroot disease in *Brassica rapa* var. *perviridis* in both chemical and organic cultivation systems. In addition, changes in the bacterial community structure in the roots of non-infected and infected plants were analyzed for both cultivation systems. To the best of our knowledge, this is the first comparative study on clubroot disease prevalence in different soil environments.

## 2. Materials and Methods

### 2.1 Preparation of chemical and organic soils

Chemical and organic soils were prepared using a base soil with chemical and organic fertilizers. Base soils were prepared by combining vermiculite, mountain soil, black soil, and peat moss in a volumetric ratio of 5:3:3:1 (v/v). To prepare the chemical soil, ammonium sulfate, calcium superphosphate, and potassium nitrate were mixed with the base soil at an N:P:K ratio of 100:100:100 (mg/kg) at weeks 0 and 2. For chemical fertilization, ammonium sulfate, calcium superphosphate, and potassium nitrate were added in ratios of 476:571:200 and 238:285:100 (mg/kg) at weeks 0 and 2 (during cultivation), respectively. The fertilization method described by previous study was employed to prepare organic soil [20]. To prepare the organic soil, organic fertilizers (cow manure 5%, oil cake 0.25%, soybean 0.25%, and bone meal 0.05%, w/w) were mixed with the same base soil at week 0. The moisture content of the soil was adjusted to 30% and maintained at 23 ℃ for one week prior to soil property analysis.

### 2.2 Analysis of soil physicochemical parameters

The levels of total carbon (TC), total nitrogen (TN), total phosphate (TP), total potassium (TK), and soil pH were analyzed prior to plant cultivation. TC was analyzed using a Total Organic Carbon Analyzer (TOC-VCPH, Shimadzu, Kyoto, Japan). TN, TP, and TK were extracted with CuSO_4_・5H_2_O, H_2_SO_4_, and H_2_O_2_ at 420 °C [21]. After extraction, TN and TP were determined using the indophenol blue method [22] and the molybdenum blue method [23], respectively. The TK in the extract was measured using an atomic absorption spectrometer (Hitachi Z2300, Tokyo, Japan). The soil pH was analyzed at a 1:2.5 ratio of soil to distilled water using a pH meter (LAQUA F-72, Horiba, Kyoto, Japan).

### 2.3 Preparation of *Plasmodiophora brassicae* spores and pathogenic soils

Resting spores of *Plasmodiophora brassicae* were obtained from infected *B. rapa* roots, and a spore suspension was prepared to create pathogenic soil via inoculation. Infected *B. rapa* roots were carefully extracted and thoroughly washed to retain only the galls. The galls were subsequently crushed in a ratio of 1:1.5 (galls to distilled water, v/w) using a mixer. The resulting suspension was filtered through 500 mm and 100 mm mesh sheets and centrifuged (500 rpm for 5 min) in order to separate the debris. The supernatant was subjected to a second centrifugation (1,000 rpm for 10 min). The precipitate containing the resting spores was suspended in distilled water. This purification process was repeated three times to ensure spore purity. A storage buffer (Hoagland solution) was added to the final precipitate to inhibit spore germination. The number of resting spores in the suspension was determined using a hemocytometer under a microscope (BX-50; Olympus, Tokyo). The spore suspension (4.0 × 10^8^ spores/g-soil) was stored at 4 °C. Chemical and organic pathogenic soils were prepared by inoculating resting spore suspensions. Three pathogenic soil samples were prepared, each with different spore concentrations (1 × 10^4^, 1 × 10^5^, and 1 × 10^6^ spores/g soil). These soils were then incubated at 23 ℃ for one week.

### 2.5 Plant cultivation and growth measurements

*Brassica rapa* var. *perviridis* were cultivated for the experiment. Seedlings of *B. rapa* were prepared for one week, and six seedlings were transplanted into each pot (3 L soil/pot). The temperature during cultivation was maintained at 23 ℃ with a light-dark cycle of 12 h light and 12 h darkness. After four weeks of cultivation, plant growth was measured in terms of fresh shoot weight.

### 2.6 Measurement of disease class and index (DI)

To assess the severity of clubroot disease, the disease class and disease index (DI) were measured. The infected roots were visually categorized into five stages in accordance with a previous study [24]: Class 0, no onset; Class 1, small galls only on the lateral roots; Class 2, small galls on the taproot; Class 3, large galls on the taproot with the lateral roots retained; Class 4, galls on the entire root. The DI was calculated for each soil treatment using the following equation:

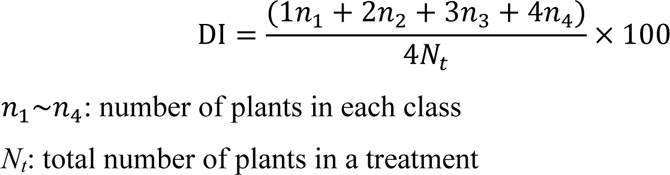

### 2.7 Investigation of root infection

Root infection in plants cultivated under both chemical and organic cultivation systems was investigated, with a spore concentration of 1×10^5^ spores/g soil. The plants were harvested on days 4, 7, 11, 14, and 28. For microscopic observation at the cellular level, the roots were thoroughly washed with distilled water, and a 1 cm hypocotyl screen from the top of the root was cut for staining. Root samples were stained by immersion in an acetic acid:ethanol (1:1) mixture for 1 min, followed by 0.01% methylene blue for 10 s [25]. After washing the stained samples, taproot tissues were placed on a glass slide for preparation and observed using an optical microscope (BX-50, Olympus, Tokyo).

### 2.8 Soil and root bacterial number analysis

The bacterial biomass of the soil was analyzed by quantifying the environmental DNA (eDNA) extracted using a slow stirring method [26]. Root bacterial biomass was also extracted using the slow stirring method with slight modifications. To extract bacterial eDNA from the root samples, the root surface was sterilized with 75% ethanol for 1 min, followed by 1% hypochlorous acid for 3 min, and washed three times with sterilized water. Then, 1 g root was crushed using a mortar and pestle for 3 min, and the pulverized root was then subjected to root bacterial eDNA extraction using the slow-stirring method. The extracted eDNA was quantified based on the intensity of the eDNA bands after electrophoresis on an agarose gel using Kodak 1D 3.6 Image Analysis Software (Kodak, CT, USA). The bacterial biomass in the soils and roots was estimated using the equation *Y* = 1.70 × 10^8^ *X* (r^2^ = 0.96), where Y and X represent the bacterial biomass g^-1^ of soils or roots and the amount of eDNA, respectively.

### 2.9 Soil and root bacterial diversity analysis by polymerase chain reaction denaturing gradient gel electrophoresis (PCR-DGGE)

The bacterial eDNA extracted from the soils and roots was used for PCR-DGGE. The 16S rRNA bacterial gene was amplified using primers DGGE-F (5’-CGCCC GCCGC GCCCC GCGCC CGTCC CGCCG CCCCC GCCCG CCTAC GGGAG GCAGCAG - 3’) and DGGE-R (5’- CCGTC AATTC CTTTG AGTTT - 3’). Further details of these methods are provided in the Supporting Information (Methods S1).

### 2.10 Metagenomic analysis of non-infected and infected root bacterial communities

Bacterial eDNA from non-infected and infected roots from chemical and organic soils was subjected to 16S rRNA sequencing for metagenomic analysis. Details of sequencing are provided in the Supporting Information (Methods S2).

### 2.11 Statistical analysis

Statistical analyses were carried out using SPSS software v. 25.0. Data were analyzed by one-way analysis of variance (ANOVA) in SPSS software (Version 19.0, Armonk, NY, USA). Tukey’s *t* test was performed to check the significance of the differences. Principal component analysis (PCA) was performed to determine the relationship between the endophytic bacterial populations and root types. Figures were processed using OriginPro software (Version 8.5, OriginLab Corporporation, Northampton, MA, USA). The data means were considered significantly different at *p* < 0.05.

## 3. Results

### 3.1 Chemical and biological properties of experimental soils

The chemical and biological properties of the experimental soils were analyzed. Chemical properties, such as TC, TN, TP, and TK, were higher in the organic soil than those in the chemical soil (Table 1). The pH values were similar in both chemical and organic soils. Bacterial biomass and diversity, which serve as indicators of biological properties, were compared between the two soil types. The bacterial biomass in the chemical soil was below the detection limit (<6.6 × 10^6^ cells/g soil), whereas the bacterial biomass in the organic soil was 3.5 × 10^8^ cells/g soil. Furthermore, the bacterial diversity in the organic soil was higher than that in the chemical soil (Fig 1). Different fertilization practices resulted in changes to the chemical and biological properties of soils.

**Table 1.** Chemical and biological properties of chemical and organic soil.

| **Soil** | **TC**<br>(mg/kg) | **TN**<br>(mg/kg) | **TP**<br>(mg/kg) | **TK**<br>(mg/kg) | **pH** | **Bacterial biomass**<br>($\times 10^8$ cells/g-soil) |
| --- | --- | --- | --- | --- | --- | --- |
| Chemical | 13,600 | 700 | 500 | 8,200 | 6.3 | n.d. |
| Organic | 41,800 | 1,300 | 1,000 | 10,700 | 6.7 | 3.5 |

\*TC, total carbon; TN, total nitrogen; TP, total phosphorus; TK, total potassium
\*n.d.: Below the detection limit ( $<6.6 \times 10^6$ cells/g-soil).

**Fig 1.**
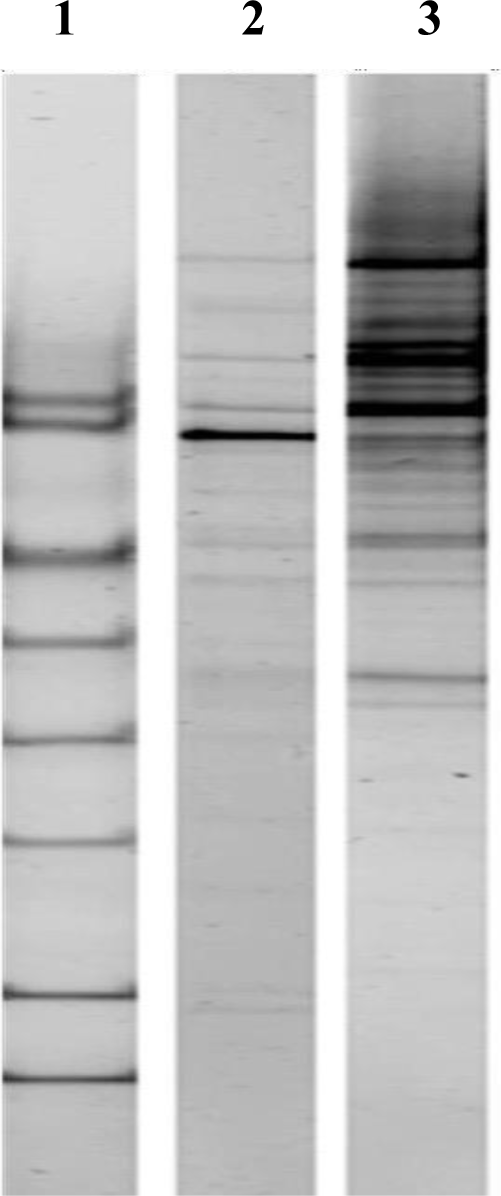
PCR-DGGE analysis of bacterial diversity in chemical and organic soils. PCR-DGGE marker (1), DNA band pattern for bacteria in chemical soil (2), and DNA band pattern for bacteria in organic soil (3).

### 3.2 Plant growth in pathogenic chemical and organic soils

*B. rapa* was cultivated in both pathogenic chemical and organic soils. Figure 2 shows the growth of *B. rapa* cultivated in chemical and organic soils with different numbers of spores. The fresh weight of *B. rapa* decreased as the spore number of *P. brassicae* increased in both chemical and organic soils. However, no significant differences in plant growth were observed when the plants were cultivated in pathogenic organic soil. (Fig. 3b).

**Fig 2.** Growth of *B. rapa* at different spore numbers in chemical and organic soils. Fresh weight in (**a**) chemical soil and (**b**) organic soil following four weeks of cultivation. Different superscript letters within columns are significantly different, as determined by Tukey’s post hoc test (*p* < 0.05).

**Fig 3.**
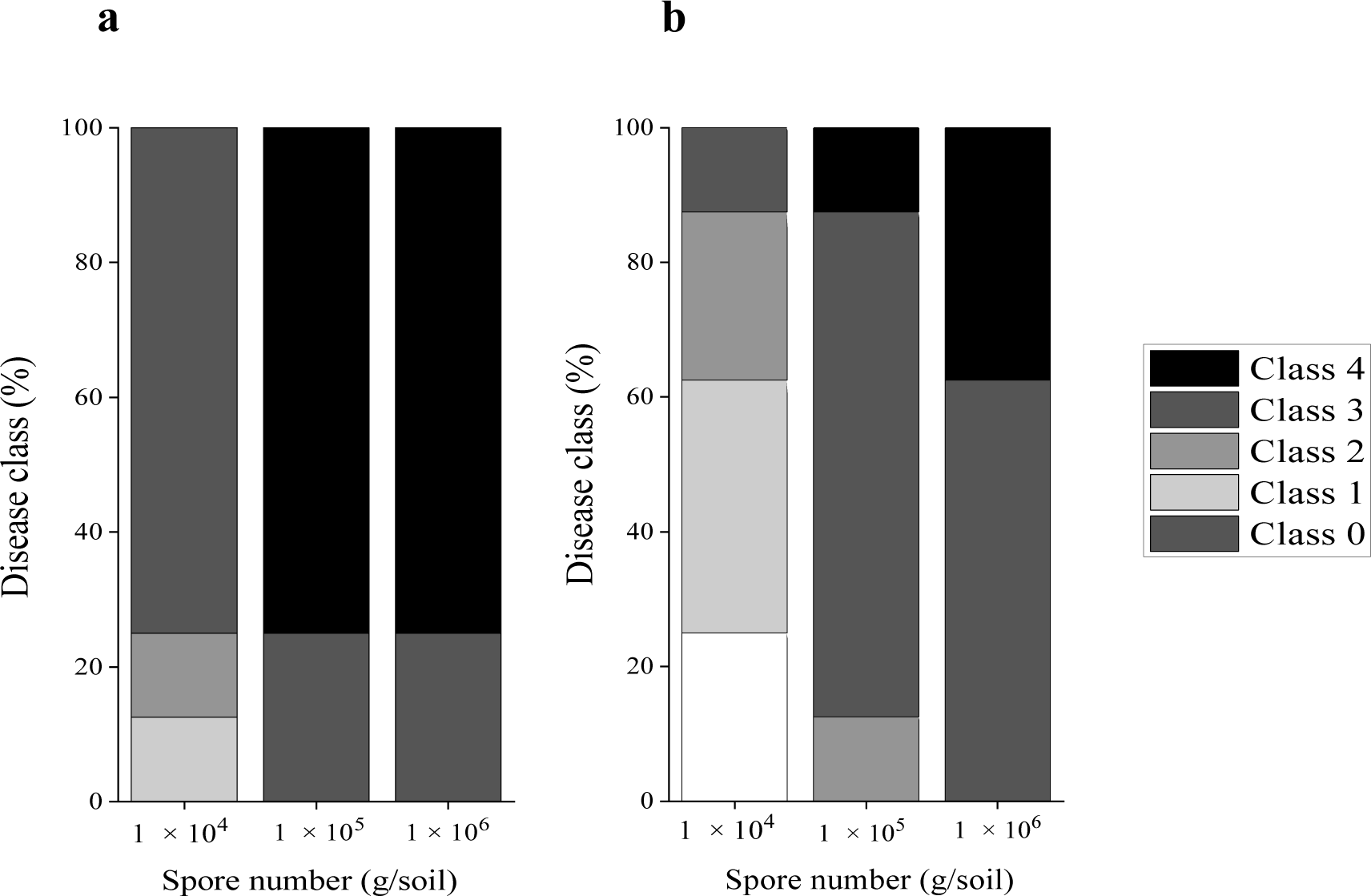
Disease class of infected roots at different spore numbers in chemical and organic soils. **a:** chemical soil, **b**: organic soil.

### 3.3 Clubroot disease class and disease index in chemical and organic soils

Clubroot disease class and DI in both chemical and organic soils were analyzed to investigate disease severity. The number of infected plants and clubroot disease classes were higher in the chemical soil than in the organic soil (Fig 3). Only non-infected (Class 0) plants were observed when the spore number was 1.0 × 10^4^ spores/g soil under organic cultivation. The number of plants with Class 4 infection was lower under organic cultivation than under chemical cultivation.

The DI of the plants was investigated at different spore numbers during chemical and organic cultivation. Higher spore numbers in the soil resulted in an increase in the DI under both cultivation conditions. However, the DI was lower under organic cultivation than under chemical cultivation (Fig 4). At a spore concentration of 1.0 × 10^4^ spores/g soil, the DI in chemical cultivation was approximately three times higher than that in organic cultivation. The same trend was observed at higher spore concentrations (1.0 × 10^5^ and 1.0 × 10^6^ spores/g soil, respectively) under both chemical and organic cultivation. These results indicate that chemical soil is more susceptible to clubroot disease infection than organic soil.

**Figure 4.**
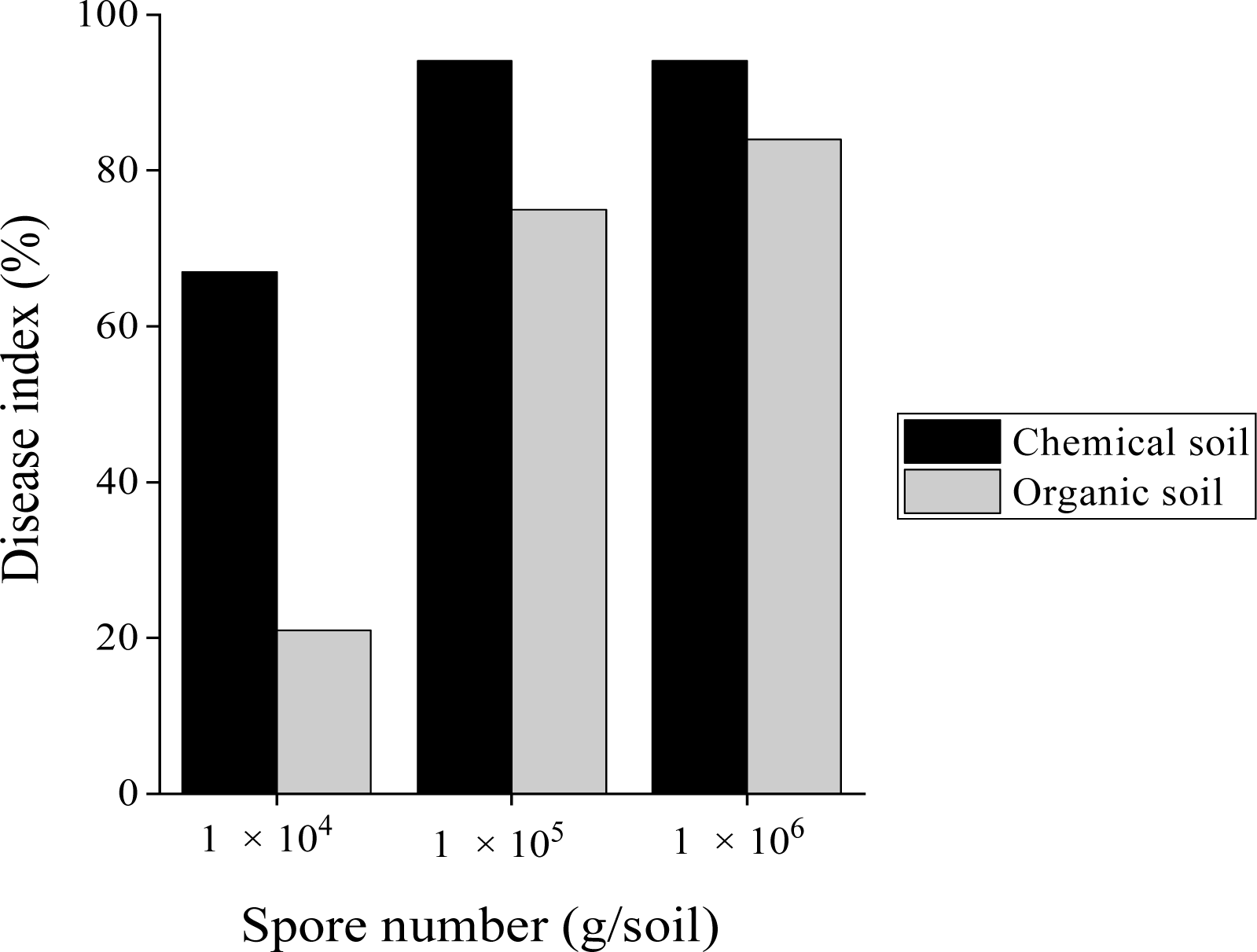
Disease index of infected plants at different spore numbers in chemical and organic soils.

### 3.4 Root infection level at different time courses

We examined root infections in both chemical and organic soils on Days 4, 7, 11, 14, and 28 of cultivation, using a spore concentration of 1 × 10^5^ spores/g soil. Root infection was observed on Days 4, 7, 11, 14, 21, and 28 of cultivation in both chemical and organic soils (Fig 5). In the chemical soil, nodules on the roots began to appear after 14 d, and lateral roots formed nodules after 28 d of cultivation. In contrast, minimal root knot formation was observed in the organic pathogenic soil after 14 d of cultivation. After 28 d of cultivation, enlarged nodules were observed; however, healthy lateral roots remained following organic cultivation.

**Fig 5.**
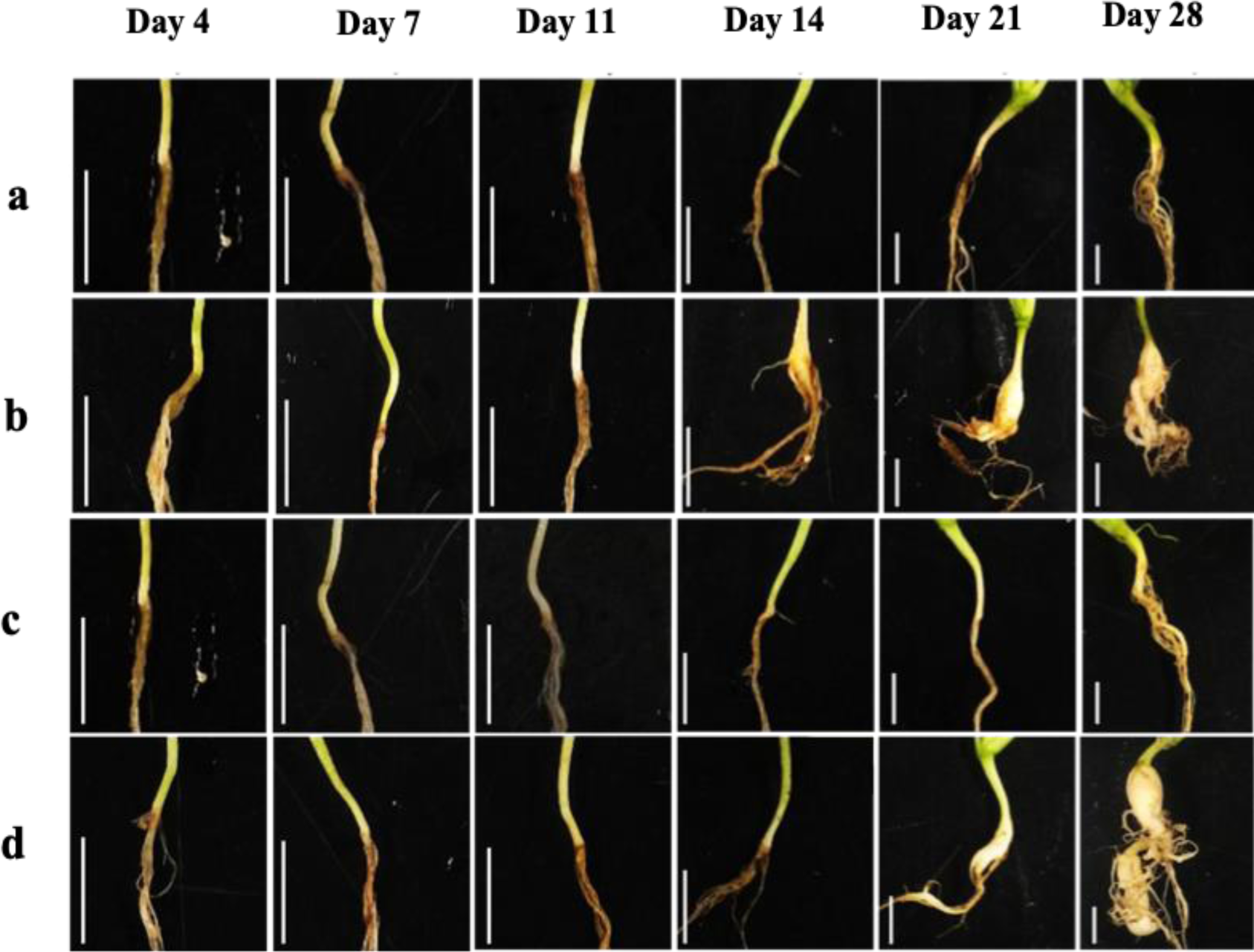
Temporal progression of root infection levels at different spore concentrations in chemical and organic soils. Root structure in (**a**) non-pathogenic chemical soil, (**b**) pathogenic chemical soil, (**c**) non-pathogenic organic soil, and (**d**) pathogenic organic soil. Scale bar: 1 cm.

The presence of intercellular spores in infected roots was investigated in both chemical and organic soils (Fig. S1). The appearance of intracellular spores in infected roots was first detected after 21 d of cultivation in both chemical and organic environments. However, the spores were more densely packed within the cells of the infected roots in the chemical soil (black area in the 28-d-old histogram in Fig. S1; RS) than in the roots in the organic soil (tissue diagram; Fig. S1) after 28 d of cultivation. The number of spores was more than four times higher in the infected roots from the chemical soil than in the infected roots from the organic soil (Table 2).

**Table 2.** Number of spores in the infected root of *B. rapa* plants after four weeks of cultivation. Data are represented as mean ± standard deviation. Different superscript letters within columns are significantly different, as determined by Tukey’s post hoc test (*p* < 0.05) (n= 3)

| Soil Type | Spore number in soils<br>(g/soil) | Spore number in roots<br>($\times 10^8$ cells/g root) |
| --- | --- | --- |
| Chemical | $1.0 \times 10^5$ | $2.10 \pm 0.21^a$ |
| Organic | $1.0 \times 10^5$ | $0.50 \pm 0.15^b$ |

Therefore, delayed nodule formation and a lower number of spores in the root cells suggest that the severity of clubroot disease was lower under organic cultivation than under chemical cultivation.

### 3.5 Bacterial biomass and diversity in roots

Bacterial biomass and diversity were analyzed in both non-infected and infected roots cultivated in chemical and organic soils. The bacterial biomass in the non-infected roots was significantly higher than that in the infected roots cultivated in both chemical and organic soils (Table 3). These results indicate that infection by the clubroot pathogen suppressed the proliferation of specific endophytic bacteria in *B. rapa*.

**Table 3.** Bacterial biomass in the non-infected and infected roots of *B. rapa* following four weeks of cultivation. Data are represented as mean ± standard deviation. Different superscript letters within columns are significantly different, as determined by Tukey’s post hoc test (*p* < 0.05) (n = 3).

| Soil type | Spore number in soil<br>(g/soil) | Disease occurrence | Bacterial biomass in root<br>($\times 10^8$ cells/g root) |
| --- | --- | --- | --- |
| Chemical | 0 | Non-infected | $7.60 \pm 3.20^a$ |
| | $1.0 \times 10^5$ | Infected | $4.40 \pm 2.10^b$ |
| Organic | 0 | Non-infected | $7.40 \pm .05^a$ |
| | $1.0 \times 10^5$ | Infected | $5.90 \pm 1.54^b$ |

PCR-DGGE was performed to analyze bacterial diversity in non-infected and infected roots cultivated in both chemical and organic soils. Bacterial diversity differed between the chemical and organic soils; however, there were similarities in the non-infected roots of both cultivation systems (Fig 6).

**Fig 6.**
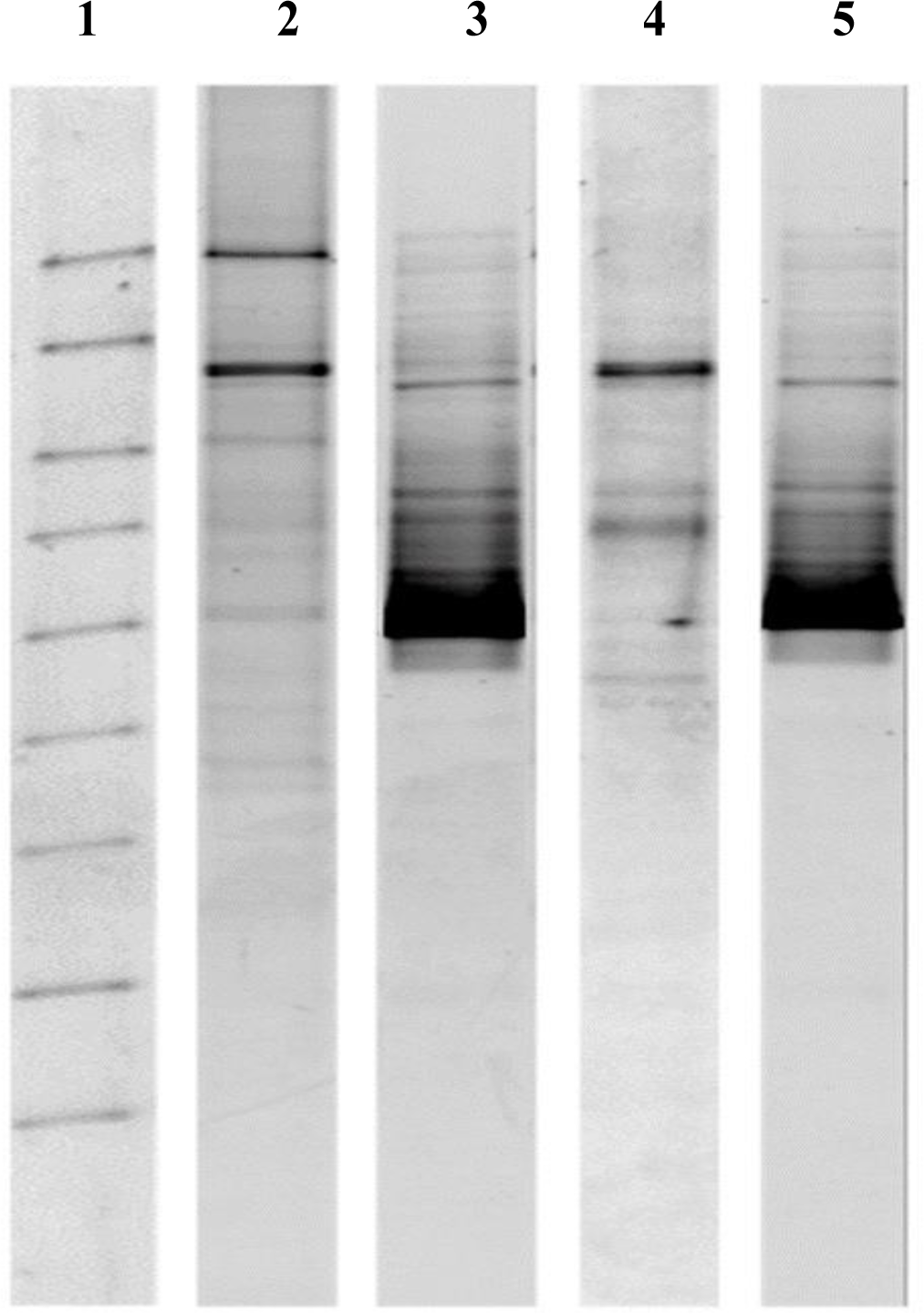
PCR-DGGE analysis of bacterial diversity in non-infected and infected roots cultivated in chemical and organic soils. (**1**) 16S PCR-DGGE markers; (**2**) DNA bands of non-infected root bacteria cultivated in chemically treated soil; (**3**) infected root bacteria cultivated in chemical soil; (**4**) non-infected root bacteria cultivated in organic soil; and (**5**) infected root bacteria of *B. rapa* cultivated in organic soil.

In contrast, the bacterial diversity in the infected roots substantially changed in both cultivation systems. Bacterial diversity, as detected by PCR-DGGE, indicated that the proliferation of specific endophytic bacteria increased following the onset of clubroot disease.

### 3.6 Changes in bacterial diversity in the infected root

Roots analyzed for bacterial diversity (i.e., PCR-DGGE) were subjected to metagenomic investigations to observe changes in the bacterial flora after infection with the clubroot pathogen. At the class level, Flavobacteria and Gammaproteobacteria were abundant in the non-infected roots of the chemically cultivated plants (Fig 7). Conversely, in the infected roots of the chemically treated soil, the number of Gammaproteobacteria increased considerably, whereas that of Flavobacteria decreased. A similar trend in bacterial numbers at the class level was observed in both infected and non-infected roots of organically cultivated plants. However, the changes in the bacterial flora of the roots differed between the chemical and organic cultivation methods. For example, the number of Betaproteobacteria decreased by 84% in infected roots compared to that in non-infected roots in chemical soil. In organic soil, it decreased by 58%. Similarly, the number of Alphaproteobacteria in the infected roots in chemical soil decreased by 95%; however, there was no change in the infected roots compared to non-infected roots in organic soil. Chloroplasts were not detected in the infected roots in chemical soil but were substantially elevated in the case of organic soil.

**Fig 7.**
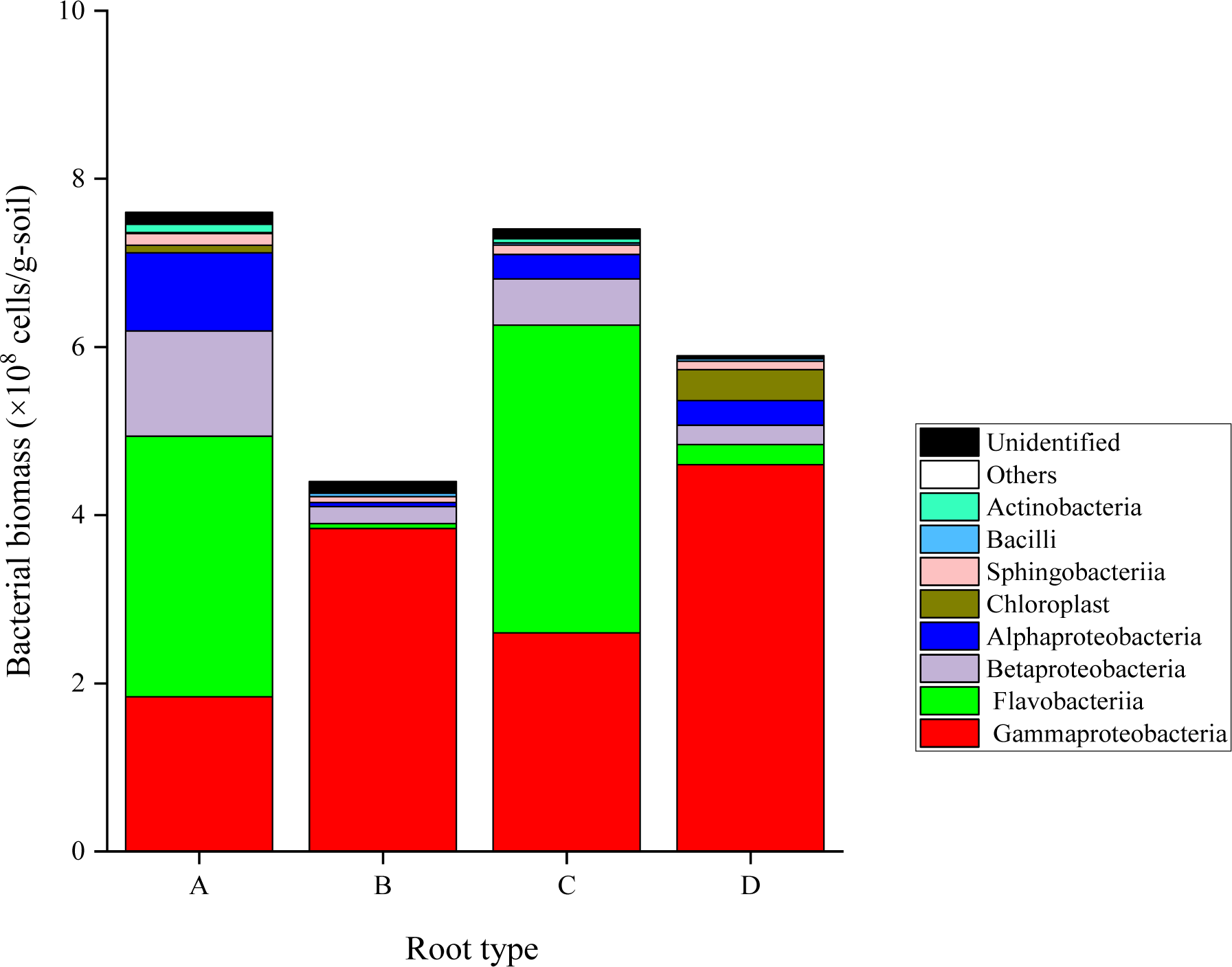
Bacterial diversity of non-infected and infected roots cultivated in chemical and organic soils. Bacterial numbers of major classes in (A) non-infected roots in chemical soil, (B) infected roots in chemical soil, (C) non-infected roots in organic soil, and (D) infected roots in organic soil.

PCA confirmed a direct relationship between the endophytic bacterial numbers at the class level and infected and non-infected roots (Fig. S2). These results indicate that the soil environment outside the roots affected the number of bacterial flora in the roots.

The relative abundance (<0.01%) at the species level was compared between non-infected and infected roots (Table S1). The relative abundance of bacterial species decreased in the infected roots in both cultivation systems. A metagenomic study of non-pathogenic and pathogenic root bacteria of *B. rapa* indicated that clubroot disease onset changes the number of bacterial flora in both chemical and organic cultivation systems.

## 4. Discussion

This study investigated the differences in clubroot disease onset in *B. rapa* cultivated in chemical and organic soils, were soil attributes differed owing to fertilization management. The DI for the occurrence of clubroot disease in organic soil was significantly lower than that in chemical soil. In addition, the disease class was lower in organic soil, with approximately 15% of plants showing no infection, even in soil with 1 × 10^4^ spores/g soil. Furthermore, the number of clubroot disease spores present in the roots in chemical soil was approximately one-fourth of that in organic soil, suggesting that clubroot disease infection was suppressed in organic soil. Bacterial biomass and diversity in organic soil were significantly higher than those in chemical soil, indicating that the abundance of soil bacteria could play a role in inhibiting clubroot disease infection. A previous study reported that soil management with improved biological properties could suppress clubroot disease [27, 28].

No significant differences in bacterial biomass within the roots were observed between the chemical and organic soils. However, the bacterial biomass in the infected roots decreased in both soils. Moreover, the bacterial diversity in the infected roots substantially changed compared to that in the non-infected roots in both soils. The significant changes in bacterial diversity in the roots infected with clubroot disease suggest that substances secreted by the clubroot pathogen affect many bacterial species in the roots. The competitive interaction between pathogens and root microbiota for nutrient availability could lead to inhibition of proliferation of endophytic bacteria.

The changes in bacterial number and diversity in the infected roots were consistent with those confirmed by previous study [29] in the endophytic microbiome of non-infected and infected *B. napus* with *P*. *brassicae*.

In the analysis of microbial communities within the roots, Flavobacteria and Betaproteobacteria were significantly reduced in infected roots, whereas Alphaproteobacteria was increased. These results indicate that specific root bacteria are affected by clubroot disease infection and that two types of bacteria, cooperative and antagonistic, are present in the roots of *B. rapa*. Flavobacteria and Betaproteobacteria, which are inhibited by the clubroot disease pathogen, may suppress clubroot disease infection. The change in specific endophytic microbiota could be attributed to the situations of microbe-microbe interaction in the infected root compartment [30]

In this study, the suppression of clubroot disease and minimal growth impairment in *B. rapa* grown in organic soil environments suggest that the presence of antagonistic bacteria, influenced by the abundance of soil bacteria and microbial diversity, contributed to the inhibition of clubroot disease infection. Maintaining or increasing soil bacterial biomass in appropriate organic soil environments could be considered a method to suppress clubroot disease infection.

## 4. Conclusions

The bacterial numbers and diversity were distinct in the chemical and organic soils. Higher bacterial numbers and diversity in the organic soil and roots *of B. rapa* root reduced the infection rate and subsequently lowered disease severity. In this study, artificial organic soil was used for the single cultivation of *B. rapa*. However, in commercial organic fields, *B. rapa* is repeatedly cultivated under the same soil conditions. Therefore, to understand the effects of continuous organic fertilization on clubroot disease severity, it is necessary to conduct experiments with repetitive cultivation using the same soil at the field scale.

## Acknowledgement

The authors thank Pitchayapa Pholkaw and Kazuyoshi MAEDA for their skillful technical assistance. The authors also thankful to Kiwako S. Araki for helpful discussion. This research work did not receive any external funding.

## Author contributions

Z. Islam conceptualization, investigation, formal analysis, writing of original draft, and data curation. M. Kubo supervision, and writing, including review and editing. TQ. Thinh investigation and writing, including review and editing.

## Data availability

The data of this study are available on request

